# Inherent population structure determines the importance of filtering parameters for reduced representation sequencing analyses

**DOI:** 10.1101/2020.11.14.383240

**Authors:** D. Selechnik, M.F. Richardson, M.K. Hess, A.S. Hess, K.G. Dodds, M. Martin, T.C. Chan, A.P.A. Cardilini, C.D.H. Sherman, R. Shine, L.A. Rollins

## Abstract

As technological advancements enhance our ability to study population genetics, we must understand how the intrinsic properties of our datasets influence the decisions we make when designing experiments. Filtering parameter thresholds, such as call rate and minimum minor allele frequency (MAF), are known to affect inferences of population structure in reduced representation sequencing (RRS) studies. However, it is unclear to what extent the impacts of these parameter choices vary across datasets. Here, we reviewed literature on filtering choices and levels of genetic differentiation across RRS studies on wild populations to highlight the diverse approaches that have been used. Next, we hypothesized that choices in filtering thresholds would have the greatest impact when analyzing datasets with low levels of genetic differentiation between populations. To test this hypothesis, we produced seven simulated RRS datasets with varying levels of population structure, and analyzed them using four different combinations of call rate and MAF. We performed the same analysis on two empirical RRS datasets (low or high population structure). Our simulated and empirical results suggest that the effects of filtering choices indeed vary based on inherent levels of differentiation: specifically, choosing stringent filtering choices was important to detect distinct populations that were slightly differentiated, but not those that were highly differentiated. As a result, experimental design and analysis choices need to consider attributes of each specific dataset. Based on our literature review and analyses, we recommend testing a range of filtering parameter choices, and presenting all results with clear justification for ultimate filtering decisions used in downstream analyses.

## Introduction

As novel technologies expand the scope of study in molecular ecology, more research is needed to fully understand how methodological and population structure inference choices can influence conclusions. Restriction site-associated DNA sequencing (RADSeq) and genotyping-by-sequencing (GBS) are examples of next-generation sequencing (NGS) technologies referred to as reduced representation sequencing (RRS). Through these methods, restriction enzymes are used to cut at specific sites throughout a genome, thereby determining which regions are sequenced (Davey & Blaxter, 2011; Elshire et al., 2011; Peterson et al., 2012; Poland & Rife, 2012). Combined with specialized downstream bioinformatics pipelines, these tools have facilitated the study of wild populations (Ellegren, 2014), including those of species without fully sequenced genomes, with a high number of SNPs (Shafer et al., 2017). RRS allows for more complete quantification of genetic diversity and differentiation, and thus more accurate inferences in population genetics and phylogenetics (Parchman et al., 2018), than do previous methods. As a result, RRS approaches have become popular in studies on conservation, invasion, and evolution (Narum et al., 2013; Parchman et al., 2018; Rius et al., 2015; Shafer et al., 2016; Shafer et al., 2015).

Within RRS, there are several techniques that vary in the type and number of restriction enzymes used, adapter ligation methods, size selection, barcoding, and type of sequence data generated (Andrews et al., 2016). In addition to the sequencing method, read depth (also called sequencing coverage: the average number of reads that align to each reference base) must be selected; higher depths provide greater accuracy of genotype calls but are costlier, so fewer individuals can be sequenced under a fixed budget. The laboratory practices best-suited for a given project depend on factors such as genome size, restriction site density, and levels of linkage disequilibrium (LD) (Lowry et al., 2017) of the organism, as well as the questions being asked (Andrews et al., 2016). While RRS approaches are useful for assessing neutral genetic variation and population structure, they may be less well-suited than other NGS technologies for studying adaptation (Lowry et al., 2017). This is because RRS does not provide information on all SNPs across a genome or transcriptome, but rather a subset of SNPs found within ‘tags’ (regions of the genome between cleavage sites of selected restriction enzymes) (Davey & Blaxter, 2011). Furthermore, because genomes are mostly composed of non-coding sequences, SNPs identified by RRS are more likely to represent neutral variation (as opposed to variation under selection) than those identified by NGS approaches such as RNA-Seq (Wang et al., 2009). Despite these issues, RRS approaches can be fine-tuned through experimental design choices (such as the type and number of restriction enzymes (Hamblin & Rabbi, 2014)) to vary fragment size selection, and through sequencing effort to increase SNP density, increasing the likelihood of capturing signals of selection (Catchen et al., 2017).

Methodological choices are not limited to the laboratory; when performing bioinformatics analyses, users can filter their data based on several parameters, each of which has a range of possible values (De Summa et al., 2017). To improve genotyping and SNP calling accuracy, data can be filtered for sequence quality, read depth, and strand bias (Nielsen et al., 2011). In RRS datasets, two common filtering parameters are the thresholds for call rate (the highest percentage of individuals in which the genotype for a locus is allowed to be missing, above which the locus will be filtered from the dataset) and minimum minor allele frequency (MAF; the lowest rate at which the less common allele of a bi-allelic locus can occur in the dataset, below which the locus will be filtered from the dataset). Varying these filtering parameters can influence which SNPs are retained for analysis, and consequently, can affect estimates of the levels of genetic diversity and population structure (Shafer et al., 2017).

Choosing a call rate threshold is challenging because retaining loci that are missing from many individuals may lead to inferences being drawn from uninformative data (Arnold et al., 2013). However, removing these loci may also skew interpretations; tags from regions with high mutation rates are the most likely to have mutations within restriction cut sites, which may prevent them from being captured (Arnold et al., 2013; Huang & Knowles, 2016). Thus, all loci in such tags may be missing from individuals that have cut site mutations, and filtering them out may cause underestimation of genetic diversity and differentiation (Huang & Knowles, 2016). Furthermore, allowing a higher call rate threshold may provide more power for population assignment due to the inclusion of more loci (Chattopadhyay et al., 2014). As a result, population inference has been shown to be affected by the percentage of missing data (Graham et al., 2020; Wright et al., 2019).

Choosing a MAF threshold also presents issues: rare alleles may suggest population expansion because their presence may reflect the introduction of new alleles to the population, and removing them may thus affect the accuracy of identifying population limits (Linck & Battey, 2017). This can also be affected by call rate filtering, particularly if there is a cut site variant that is specific to a population. Researchers remain unsure of the best practices in selecting values for these parameters, other than to perform analyses multiple times, including or excluding samples or loci with high rates of missing data to see if results are consistent (Grünwald et al., 2017). In cases where the results appear to differ based on the filtering steps, selection of the appropriate filtering thresholds may be important for accurate ecological and evolutionary interpretations, but it is difficult to predict which answers are correct.

Basic strategies for bioinformatics analysis of RRS data exist. For example, many potential issues with datasets can arise during library preparation or locus reconstruction; these should be identified through quality control measures and mitigated with the appropriate filtering steps (O’Leary et al., 2018). Calculating basic per-locus statistics provides information on issues such as spurious allele calls, private alleles, fixed alleles, and missing data (Grünwald et al., 2017). When characterizing population structure, results should be cross-validated by performing both model-based and nonparametric methods; model-based approaches have traditionally been used, but are more error-prone than nonparametric approaches (Linck & Battey, 2017). One such error is the confounding of population structure inference by SNPs found in only one individual (singletons), which should be filtered out (Linck & Battey, 2017).

Although best practice recommendations for RRS bioinformatics are emerging, it is unclear how often they are followed. Furthermore, additional recommendations may be necessary; we know that filtering thresholds can affect results, but we do not know if this is true under all conditions, or how the magnitude of these effects may change depending on biological variations (i.e. heterozygosity, partial ploidy, genome size, repeats) that alter the intrinsic properties of a dataset. Here, we investigate the range and consistency of parameter choices in RRS studies, and test the effects of filtering thresholds (call rate threshold and MAF) on datasets that vary in their inherent level of genetic differentiation between populations. To address this, we: (1) performed a literature review documenting the methods (particularly filtering choices) and levels of genetic differentiation and diversity reported among 209 RRS studies, and (2) analyzed seven simulated GBS datasets and two empirical datasets with varying levels of population structure and read depths. From the analysis of our simulated data, we predicted (1) that the effects of filtering choices would be most pronounced in the dataset with the least population structure, and would decline in severity in datasets with increasing population structure, and (2) that the effects of filtering choices would be more pronounced in datasets with lower read depths.

## Methods

### Literature review of RRS studies on wild populations

We compiled RRS studies by performing a Topic Search (TS) on Web of Science, which searches for the provided terms in titles, abstracts, and keywords within all available records. Our search term was “TS=((“reduced representation sequencing” OR “RADseq” OR “ddRAD” OR “GBS” OR “epi-GBS” OR “genotyping by sequencing”) AND “population gen*”)” and included papers up until mid-2020 (last accessed 7 July 2020). We manually filtered the results (N=500) to find relevant studies that performed population genetic analyses using RRS data. We eliminated those that did not assess genetic structure in wild animal populations (e.g. captive studies, domestic animals, human disease, plants). This yielded a total of 209 papers, from which we extracted information on study taxa, sample size, sequencing methods, bioinformatics pipelines, filtering parameter choices, number of resulting reads and SNPs, number of genetic groups, measures of genetic differentiation and diversity, and availability of supplemental information and raw data.

### Simulation of GBS datasets with varied population structure

We used GBSSim v0.5 (Hess et al., 2018; https://github.com/anshess/GBSSim), a package written in Julia (Bezanson et al., 2017), to create seven GBS scenarios, manipulating the population history events of each to produce varying levels of population structure (Figure 1A, Table S1). We simulated all scenarios to comprise four subpopulations of increasing relatedness (Figure 1B-C). The code outputs included a ‘true genotypes’ matrix and a zipped FASTQ file (requiring processing and filtering) for each dataset. We replicated each of the seven population history scenarios five times; the FASTQ files were further simulated at two different mean read depths (depth = 6 and depth = 25). In total, this resulted in 35 true genotypes matrices (7 scenarios × 5 replicates) and 70 zipped FASTQ files (7 scenarios × 5 replicates × 2 read depths).

**Figure 1.**
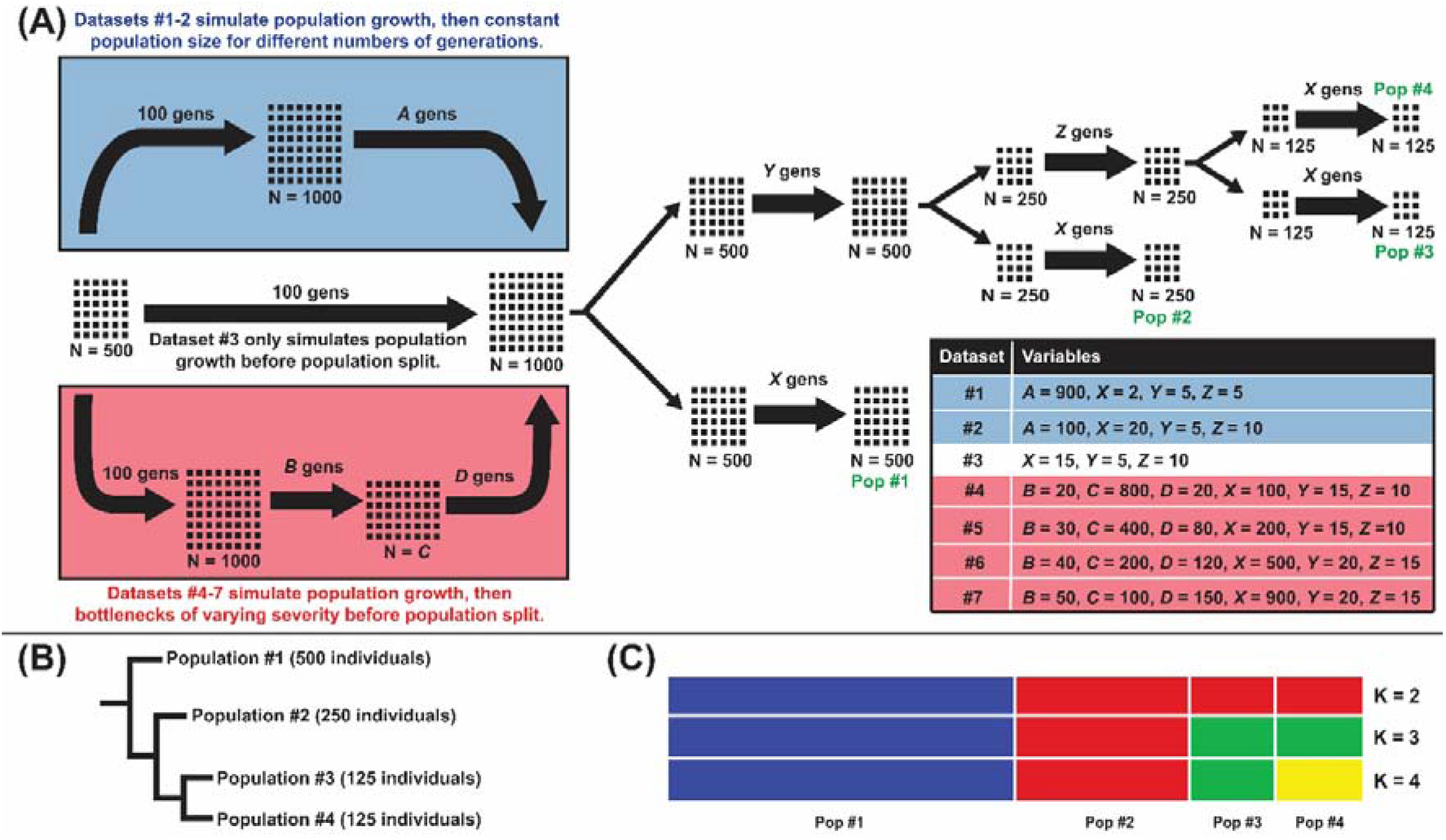
(A) Population history events of seven simulated datasets with varying degrees of structure. (B) Cladogram showing three splits occurring in the population history of all simulated datasets. (C) Expected population structure results of all simulated datasets at K=2,3,4.

Based on our literature review, we estimated that the median vertebrate genome size in these studies is 2,280 centimorgans (cM) and the average unfiltered SNP density is approximately 26 kilobases per SNP. Due to constraints with computational run times, our simulated genome size was 100 cM across one chromosome; for consistency with the average SNP density from our literature review, we simulated 4,000 SNPs before filtering in each of our datasets.

### Acquisition of empirical RAD datasets with high and low structure

To explore whether the findings of our simulation analyses extend to data obtained from wild populations, we downloaded raw reads of two publicly available ddRAD datasets from the NCBI Sequence Read Archive (SRA). The first dataset (accession: PRJNA328156) (Trumbo et al., 2016), which we call the “empirical low” dataset, has relatively low read depth and low population structure in Australian cane toads (*Rhinella marina*). We subset this dataset to isolate two populations: one in eastern Queensland (QLD; N=179), and the other in the western Northern Territory (NT; N=441). The second dataset (accession: PRJNA268025) (Bell et al., 2015), which we call the “empirical high” dataset, demonstrates relatively high read depth and high population structure in African reed frogs (*Hyperolius molleri*). We also subset this dataset to isolate two populations: one on the island of Príncipe (N=17), and the other on the island of São Tomé (NT; N=54).

### Processing of reads, SNP calling, and filtering

We used Stacks v2.0 (Catchen et al., 2011) to process all RRS data (code available on GitHub via hyperlink). First, we used the process_radtags program to remove low quality reads from the FASTQ files using the program’s default parameters (quality score of 10), truncating reads to a final length of 60 bases. Next, we used the denovo_map pipeline to perform a *de novo* assembly for each sample, align matching DNA regions across samples (called ‘stacks’), and call SNPs using a maximum likelihood framework. We then filtered the results using the populations program, using one random SNP per locus and four combinations of parameter choices (MAF=0.05, call rate threshold = 0.5; MAF = 0.05, call rate threshold = 0.2; MAF = 0.01, call rate threshold = 0.5; MAF = 0.01, call rate threshold = 0.2). The results were written to a VCF file and a STRUCTURE file, which could be read into fastStructure (Raj et al., 2014). We recorded the number of SNPs and mean read depth of each dataset after filtering.

### Evaluation of genetic differentiation and diversity

To quantify levels of genetic differentiation and diversity, we computed basic statistics in the hierfstat (Goudet, 2005) package in R (Team, 2016). We calculated global F_ST_ (mean across all loci), pairwise F_ST_ (between the four simulated genetic groups), and expected heterozygosity (He). We also computed 95% confidence intervals (CIs) for all pairwise F_ST_ values using the bootstrapping method (number of bootstraps = 100 across loci) performed by the StAMPP package (Pembleton et al., 2013). All statistics were calculated for every replicate of every dataset.

### Inference of population structure

We used fastStructure to infer population structure from both the simulated and empirical datasets using a variational Bayesian framework for calculating posterior distributions, and to identify the number of genetic clusters in our dataset (K) using heuristic scores (Raj et al., 2014). We ran fastStructure with a simple prior ten times (K= 1 to 10, a range of values around the true value of K = 4) for every replicate of every dataset. We then generated a consensus structure plot from the meanQ files of the five replicates of each dataset using CLUMPAK (Kopelman et al., 2015).

To see if inferences of population structure were consistent across approaches (model-based versus non-parametric methods), we also used adegenet (Jombart & Ahmed, 2011) to perform discriminant analysis of principal components (DAPC) on all datasets. DAPC is a multivariate approach that identifies the number of genetic clusters using K-means of principal components and a Bayesian framework (Jombart & Ahmed, 2011). Our first input for DAPC was a set of observed genotypes for every locus in each dataset. Our second input for DAPC was a set of allelic dosages (AD) for every locus in each dataset; to calculate this, we first obtained the genotype probabilities of every locus using the KGD package (Dodds et al., 2015). We then converted genotype probabilities to allelic dosages as follows:

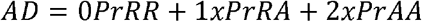

where R is the reference allele and A is the alternate allele. We determined the number of principal components (PCs) using the xvalDapc tool in adegenet, which tests ranges of PCs and identifies the optimal value (Jombart & Ahmed, 2011). Please see data accessibility statement for data and code.

## Results

### Literature review: Parameter choices made in RRS studies on wild populations

Our literature search (Supplemental File I) revealed a high degree of variation across studies in thresholds of call rate (0-0.98) and MAF (0.0038-0.25), as well as in minimum Phred scores (5-40) and read depths (1-50). We found that only 21% of studies ran multiple combinations of filtering parameter choices to check the consistency of the results, 3% cited recommendations from other papers, and 68% did not provide justification for their choices. Although many different call rate thresholds were selected across the 209 studies, values for most studies fell between 0.11-0.20 (Figure 2A). Selections of MAF were more consistent, and 0.05 was the most common choice (Figure 2B). There was a wide range of sample sizes across studies (13-3234) and approximately 56% of studies removed samples for reasons such as not passing quality filters. Approximately 66% of studies assessed population structure using both model-based (STRUCTURE, ADMIXTURE) and non-parametric (PCA, DAPC) methods. Reporting of measures such as global and pairwise F_ST_, F_IS_, and expected and observed heterozygosity (He and Ho, respectively), was inconsistent (Figure 2C). In 73% of studies, an accession number was provided linked to databases such as the SRA or Dryad, where raw sequence files or downstream files could be downloaded.

**Figure 2.**
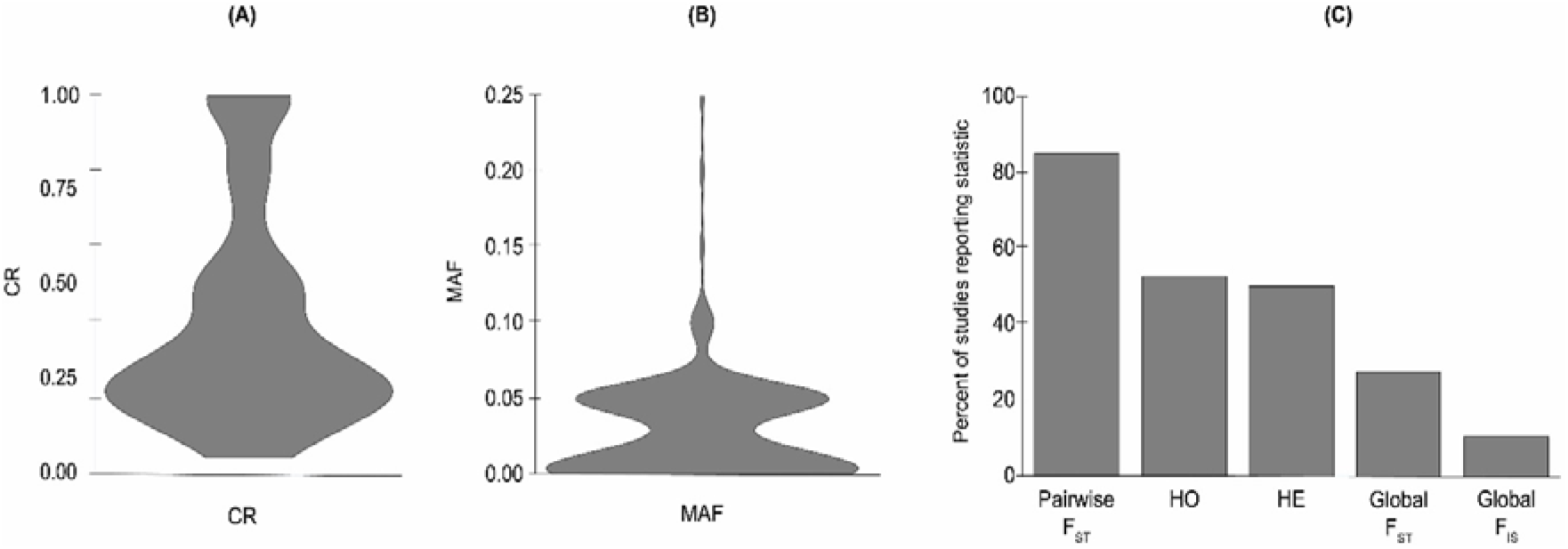
Violin plot showing frequency of choices in (A) call rate threshold values and (B) minimum minor allele frequency (MAF) values when filtering reduced representation sequence (RRS) data. A literature review was performed on 209 RRS studies, and choices of call rate threshold and MAF were extracted from each study (Table 1; full table in Supplemental File 1). (C) Percentages of 209 reduced representation sequencing (RRS) studies that report each of five population genetics statistics. A literature review was performed, and reports of global F_IS_, global F_ST_, pairwise F_ST_, expected heterozygosity (H_E_), and observed heterozygosity (H_O_) were extracted from each study (Table 1; full table in Supplemental File 1).

### Experimental determination of effects of filtering choices on inference of population structure

Our methods for assessing population structure were designed for datasets with genotypes without errors, and little missing data. Thus, one caveat of our results is that different filtering choices may be optimal if approaches that accommodate the errors are used (Bilton et al., 2018a; Bilton et al., 2018b). For this reason, we used two methods to assess population structure of our simulated data: a model-based approach implemented in fastStructure and a non-parametric approach implemented in DAPC. DAPC was performed with two types of input: observed genotypes, and allelic dosages.

To test the impacts of filtering on inferences of population structure, we filtered the datasets through every combination of two different call rate threshold values (more stringent = 0.2, less stringent = 0.5) and two different MAF values (more stringent = 0.05, less stringent = 0.01). When using the model-based fastStructure, consistency with the results from using true genotypes depended on the interaction between read depth, filtering choices, and inherent levels of differentiation in the dataset (Figure 3A-B). However, it should be noted that true genotypes were unfiltered.

**Figure 3.**
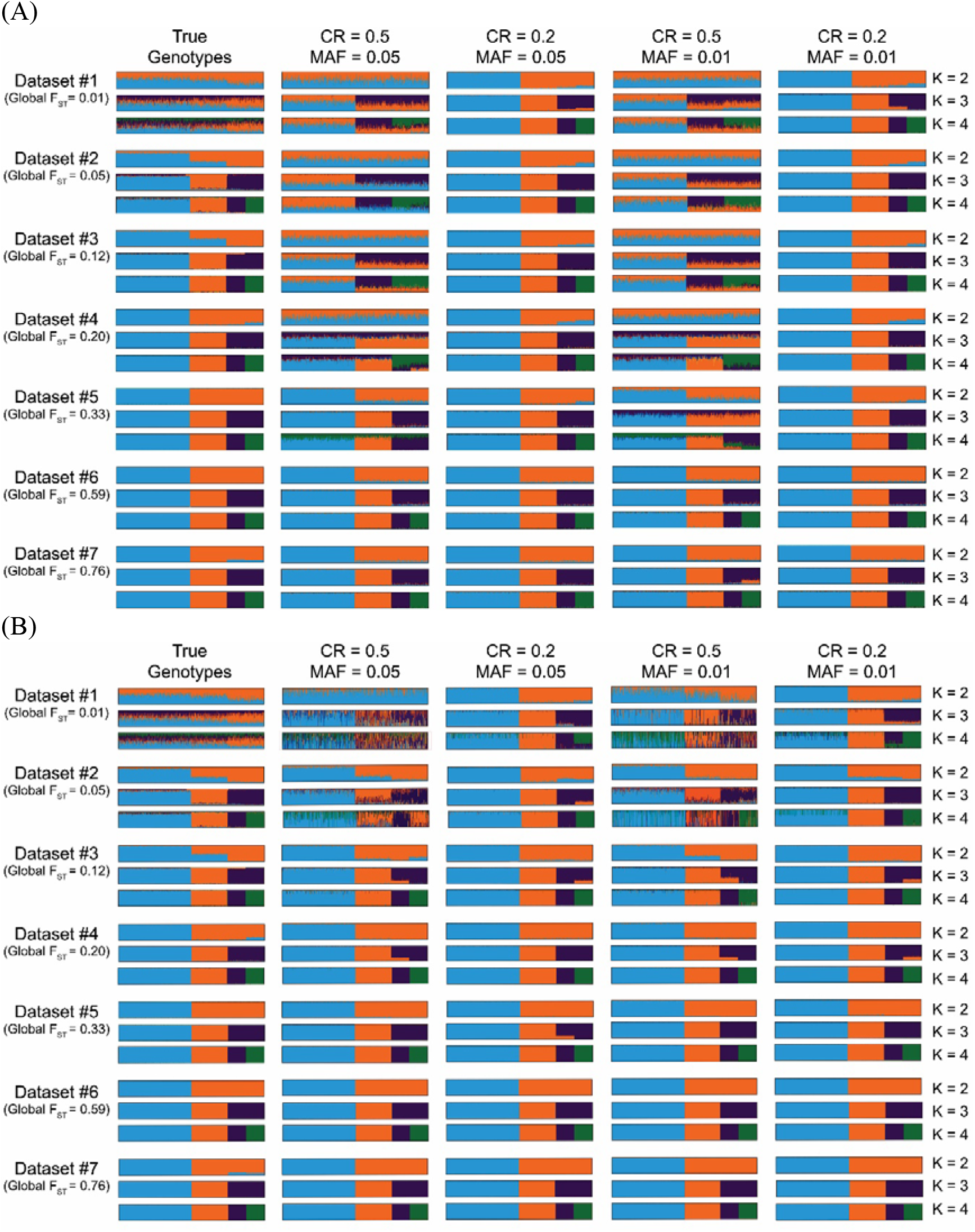
Genetic structure in seven simulated GBS datasets with varying levels of population differentiation and (A) mean read depth = 6, or (B) mean read depth = 25. All structure plots represent a consensus image of five different replicates for each dataset. Models were run four times with different combinations of filtering parameter choices (minimum minor allele frequency and call rate threshold). Results are shown for the model at K=2,3,4. Global F_ST_ values are estimated from true genotypes data. Individuals are plotted in population order.

Generally, more stringent filtering (particularly with MAF) led to a higher mean depth and lower number of SNPs retained in each dataset (Supplemental File II). With a mean read depth of 25 (as opposed to 6), the disparity between filtering choices was much smaller, and more SNPs were retained overall.

In dataset #1 (low differentiation), population differentiation was so subtle that the true genotypes did not distinguish the four populations. Across both mean read depths (Figure 3A-B), the less stringent call rate threshold choice produced results that were most consistent with those of the true genotypes (little population division). Interestingly, the more stringent call rate threshold choice allowed for detection of the four populations; although this was not consistent with the results from the true genotypes, this may simply be because the true genotypes were not filtered.

In datasets #2-4 (slightly higher differentiation than dataset #1), the four populations were distinguished by the true genotypes. At a mean read depth of 6 (Figure 3A), these four populations were only detected when using the more stringent call rate threshold choice, and structure was otherwise unclear at K=2,3,4 (we knew that K=4 was the correct answer, but also had expectations about how populations would cluster together at K=2 and K=3); choice of MAF did not appear to affect the results. At a mean read depth of 25 (Figure 3B), the four populations were detected regardless of filtering choices, and all expectations were met at K=2,3,4.

In dataset #5 (moderately differentiated), with a mean read depth of 6 (Figure 3A), population structure was clearer than in datasets #1-4 (particularly when using the more stringent choice of MAF), but the four populations were still only detected when using the more stringent choice of call rate threshold. With a mean read depth of 25 (Figure 3B), the four populations were detected regardless of filtering choices. In datasets #6-7 (highly differentiated), the four populations were detected regardless of mean read depth and filtering choices.

The findings from our simulations were corroborated by those of the empirical datasets. In the empirical low dataset, which has relatively low read depth and inherent levels of genetic differentiation comparable to simulated datasets #2-3, the two populations were only detected when using the stricter choice of call rate threshold (Figure 4). In the empirical highly differentiated dataset, which has relatively high read depth and inherent levels of genetic differentiation comparable to datasets #5-6, the two populations were detected regardless of filtering choices (Figure 4).

**Figure 4.**
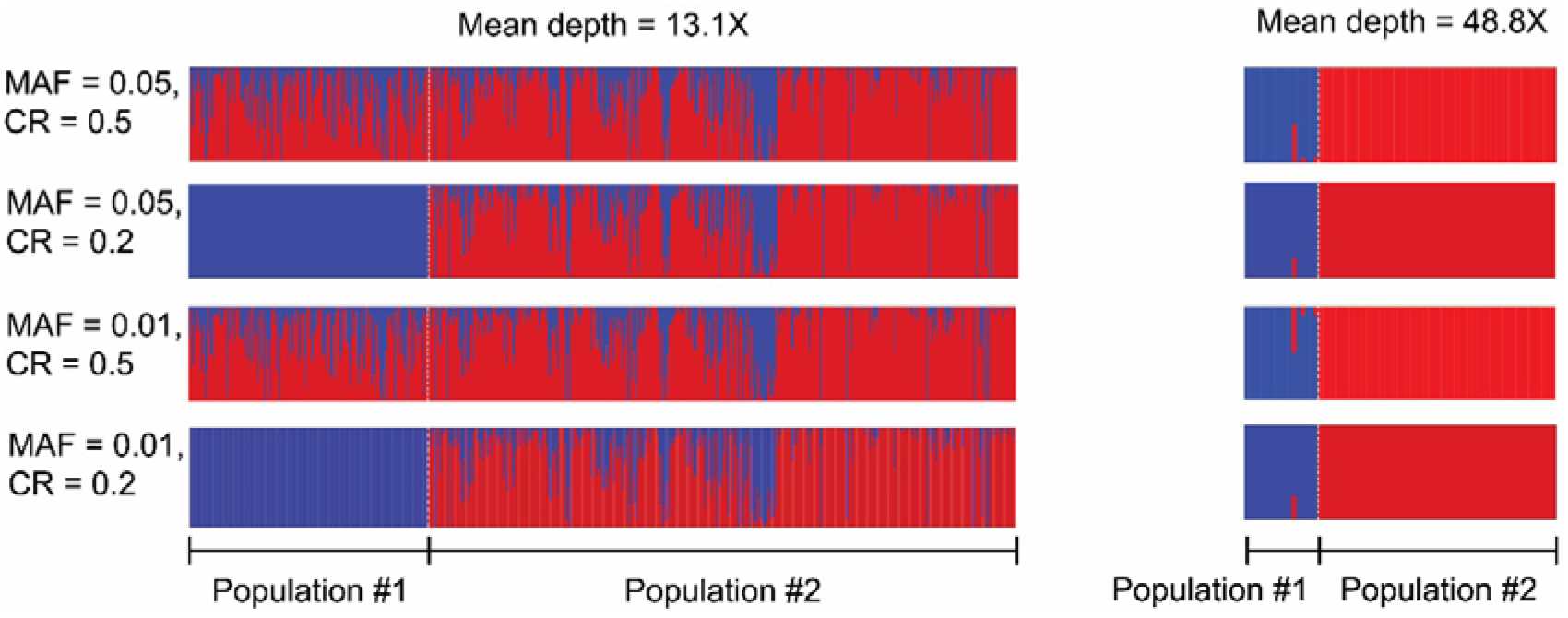
Genetic structure of two populations in each of two empirical ddRAD datasets with varying levels of differentiation and read depths. Models were run four times with different combinations of filtering parameter choices (minimum minor allele frequency and call rate threshold). Results are shown for the model at K=2.

### Non-parametric inference of population structure

When using the non-model-based DAPC, there were no methods to determine a consensus of all replicates. However, because the replicates were consistent within datasets and read depths when using observed genotypes (Figure S1A-B) or allelic dosages (Figure S2A-B), we performed DAPC on the third replicate of each dataset. In all datasets, the DAPC results were consistent whether observed genotypes (Figure 5A-B) or allelic dosages (Figure S3A-B) were used.

**Figure 5.**
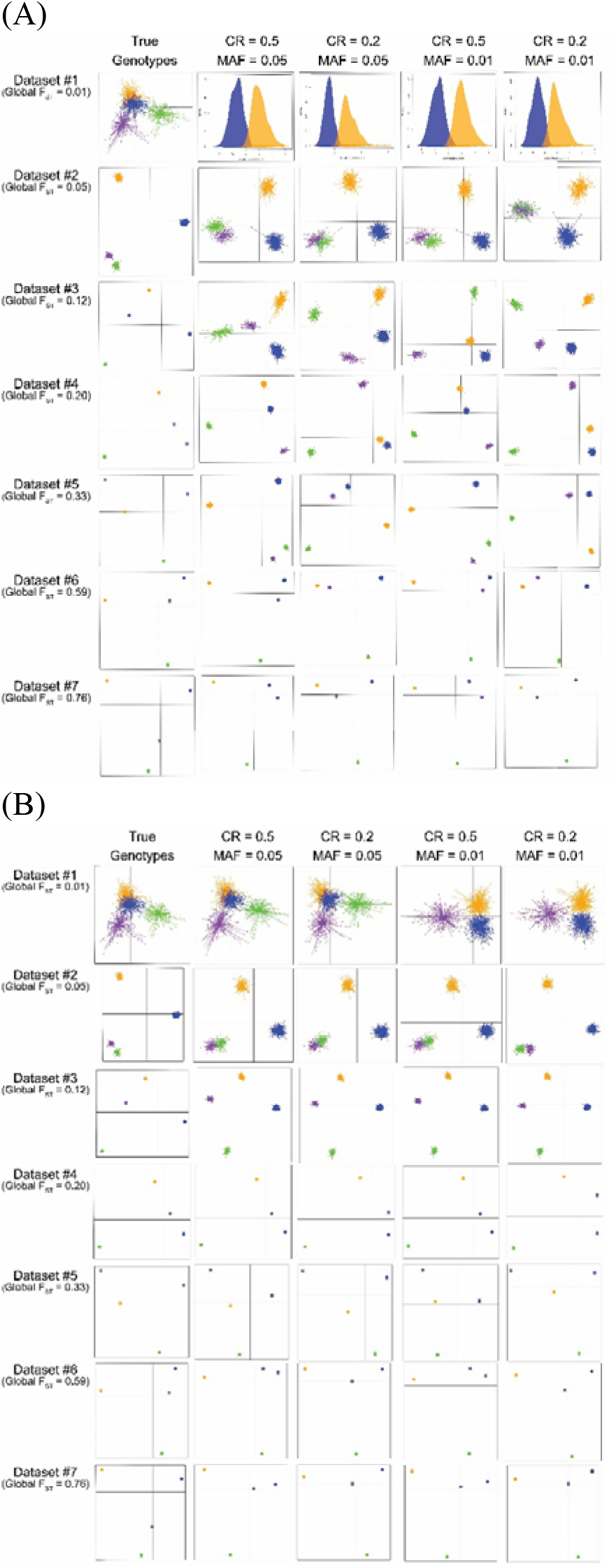
Discriminant analysis of principal components (DAPC) plots of four populations in one replicate of each of seven simulated GBS datasets with varying levels of differentiation. These were constructed using observed genotypes for every locus in each dataset. PCAs were run four times with different combinations of filtering parameter choices: minimum minor allele frequency (MAF) and call rate threshold. Read depth was also varied; (A) mean depth = 6 and (B) mean depth = 25. When DAPC detects only two populations (K=2), only a single discriminant function is retained, and densities of individuals on that function are plotted.

Unlike the fastStructure method, DAPC on the true genotypes of dataset #1 distinguished the four populations. With a mean read depth of 6 (Figure 5A), the results for dataset #1 were inconsistent with the true genotypes, regardless of filtering choices. With a mean read depth of 25 (Figure 5B), the results were closer, but they were only correct when using the more stringent MAF choice. In datasets #2-7, the results of the DAPC were mostly consistent with the true genotypes regardless of filtering choices at both read depths. Choosing the more stringent MAF choice seemed to improve accuracy in datasets with lower levels of population differentiation (datasets #2-3) with a mean depth of 6, but was not necessary at higher levels (datasets #5-7) or with a mean depth of 25. The results were generally more consistent with the true genotypes when using a read depth of 25, and this disparity was greater at lower levels of population differentiation (datasets #2-4).

Once again, the findings of the empirical datasets supported those of the simulated datasets. In both empirical datasets, detection of the two populations was consistent across filtering choices and read depths (Figure 6).

**Figure 6.**
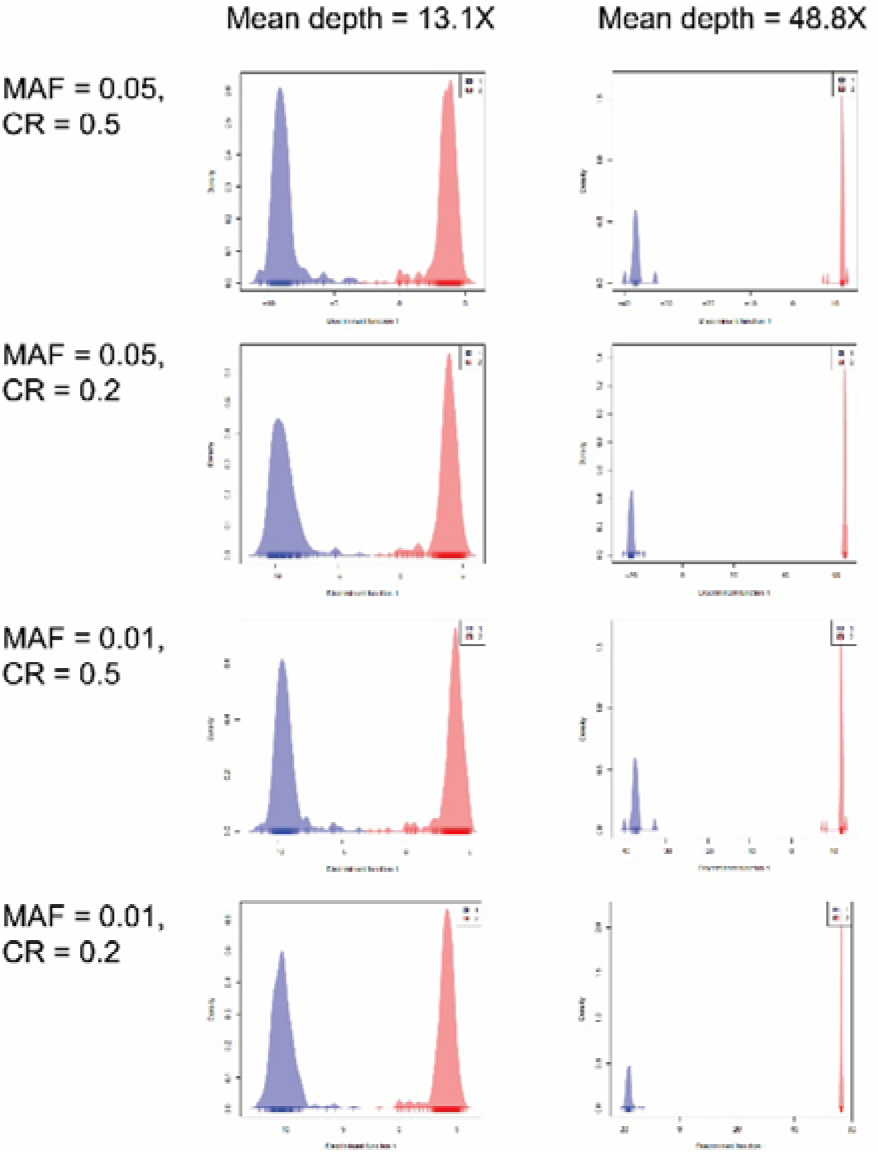
Discriminant analysis of principal components (DAPC) plot of two populations in each of two empirical ddRAD datasets with varying levels of differentiation. PCAs were run four times with different combinations of filtering parameter choices (minimum minor allele frequency and call rate threshold.

### Experimental determination of effects of filtering choices on calculations of genetic differentiation and diversity

To test the impacts of filtering on estimations of genetic differentiation and diversity, we filtered our simulated and empirical datasets through every combination of two different call rate threshold values (more stringent = 0.2, less stringent = 0.5) and two different MAF values (more stringent = 0.05, less stringent = 0.01). Estimations of global and pairwise F_ST_ in the simulated datasets were generally consistent with the true genotypes regardless of filtering choices (Table 2, full results in Supplemental File II). At a mean depth of 6, F_ST_ was slightly underestimated in the lower differentiation datasets (#1-4), but slightly overestimated in the higher differentiation datasets (#5-7; Figure S4A); all estimations of F_ST_ were more consistent with estimates from the true genotypes at a mean depth of 25 (mean error, depth 6 = −2.37%; mean error, depth 25 = −0.08%; t=-2.25; p=0.015). Similarly, estimations of global and pairwise F_ST_ were consistent across filtering choices in both empirical datasets (Table 2, full results in Tables S4-S5). Pairwise F_ST_ CIs were wider when using the more stringent choice of call rate threshold (but not MAF) at a mean read depth of 6, but because these CIs also became narrower as inherent genetic differentiation increased, these effects were less pronounced in the datasets with higher population structure (Supplemental File II). This disparity did not occur with a mean depth of 25, which also had a low percentage error (mean error, depth 6 = −3.13%; mean error, depth 25 = −0.21%; t=-2.87, p=0.003).

**Table 2.**
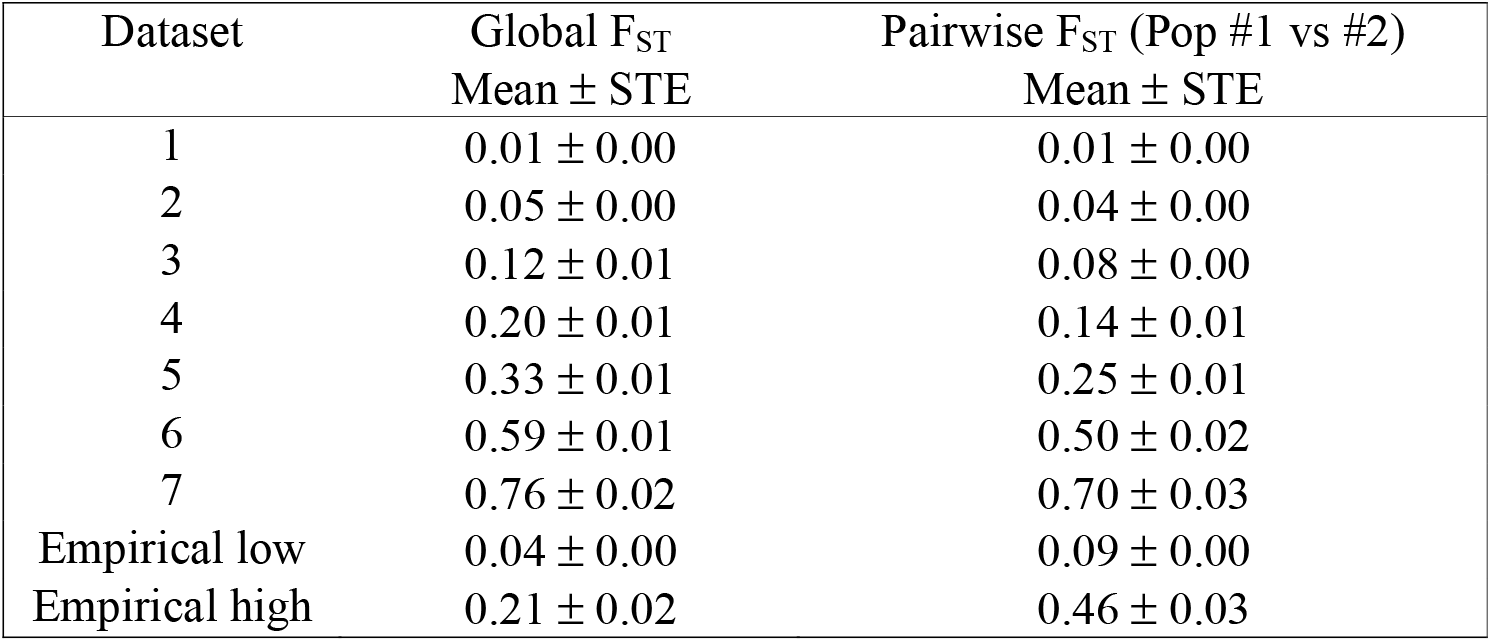
Global and pairwise F_ST_ with standard error (STE) in seven simulated GBS datasets and two empirical ddRAD datasets with varying levels of population structure. For the simulated datasets, these values were calculated from the true genotypes and averaged across five replicates of each dataset. Pairwise F_ST_ values are displayed between the first two populations (of four total) in each simulated dataset, and between the two genetic clusters in each empirical dataset. The full set of global and pairwise F_ST_ values, as well as the expected heterozygosity of each population, are available in Supplemental File II (simulated datasets) and Tables S2-S3 (empirical datasets).

Using the more stringent choices of call rate threshold and MAF generally produced higher values of He in all simulated and empirical datasets, except for the dataset with the highest structure (#7) at read depth 25 (Supplemental File II & Table S3). Interestingly, at a mean depth of 6, using the more stringent MAF choice generally produced inflated He estimates compared to true genotypes, while using the less stringent MAF resulted in decreased values of He compared to true genotypes (Figure S4B). Using more stringent MAF filtering produced more accurate He results in datasets with lower levels of differentiation (#1-3), but this did not occur in datasets with intermediate levels of differentiation (#4-6). Further, in the dataset with the highest leves of population differentiation (#7), all parameter sets from the filtering depth of 6 produced accurate values of He. At a mean depth of 25, patterns were similar to those from filtering depth of 6, except that all He values were higher in the former. Using the less stringent MAF values consistently produced more accurate results, except in the dataset with the highest level of population differentiation (#7), where all estimates of He were accurate.

## Discussion

The results of our literature review demonstrate high variation across RRS studies in experimental design choices. Only 24% of the studies we reviewed explained the rationale for their filtering choices (e.g. trialing a range of options or referencing other studies). This is understandable because considerations for ‘best practices’ when analyzing NGS data are still developing. However, unjustified choices of filtering thresholds may produce erroneous results because parameter choices can influence downstream analyses (Huang & Knowles, 2016; Linck & Battey, 2017).

We hypothesized that the extent to which filtering choices affect inference of population structure depends on the inherent levels of genetic differentiation in a dataset, with more highly differentiated populations being less affected due to their more pronounced differences. To test this hypothesis, we created seven GBS datasets with varying levels of genetic differentiation at two different read depths, and then analyzed these datasets using every combination of two different call rate threshold values (more stringent = 0.2, less stringent = 0.5) and two different MAF values (more stringent = 0.05, less stringent = 0.01). With a low mean read depth of 6, the fastStructure results of our simulations support our hypothesis; in the datasets with low population structure, stringency with call rate threshold (i.e. low levels of missing data) is required to detect separate populations. This was not the case in datasets with higher differentiation, suggesting that stricter filtering of call rate threshold is important for detection of finer scale population differentiation. In dataset #5, with intermediate to high levels of differentiation, we begin to see clearer results while using the less stringent call rate threshold, and the populations are more accurately detected when using the stricter MAF threshold than when using the less strict choice. This result suggests that choice of MAF affects the results in similar ways to call rate threshold, but that the effects of MAF are masked by the much stronger effects of call rate threshold in datasets with very low levels of differentiation. At sufficiently high levels of differentiation to reduce the effects of call rate threshold, however, we can see the effects of MAF choice as well. Both filtering parameters are likely important for filtering out noise that obscures differences between populations in the dataset. However, their effects are smaller when using the higher mean read depth (25). This is likely due to the retention of a greater number of high-quality genotypes in the dataset, allowing for more accurate inference of population structure. Interestingly, these effects are also much weaker when using DAPC, further reinforcing suggestions to use both model-based and non-parametric methods.

In dataset #1, which has the lowest levels of differentiation (F_ST_ < 0.01), the four populations are not distinguished by the true genotypes in fastStructure; however, using the more stringent call rate threshold choice when filtering clearly delineates these four populations, over-accentuating true population structure. The less stringent call rate threshold produces results that are closer to those of the true genotypes‥ However, the four populations are distinguished by the true genotypes in DAPC, and using the more stringent MAF choice at a higher read depth provides the most accurate results. This suggests that different filtering choices may be optimal for the same dataset depending on the analyses being performed.

In all other datasets, which have higher levels of differentiation (F_ST_ = 0.05-0.76) than dataset #1, the population structure results of all filtering parameter choices or read depths indicate the correct number of populations. This is true for both fastStructure and DAPC analyses, but less stringent call rate filtering results in less definitive results in fastStructure analyses. We note that our simulated datasets do not incorporate the non-random distribution of missing data across genetic groups (such as cut site mutations and structural variation that differ between populations due to genetic drift), where the amount of missing data is likely proportional to genetic distance (Huang & Knowles, 2016), but assumes that captured loci only provide single nucleotide variation. This is sometimes seen in empirical data, and in such cases, high stringency with a call rate threshold may be counterproductive. This is because removing informative loci that are not sequenced in some or most individuals may bias the data by only retaining loci with low mutation rates, potentially obscuring differences among genetically distinct groups (Huang & Knowles, 2016). Discerning if or when this arises is largely dependent on individual dataset characteristics, but further highlights the importance of testing and reporting a range of filtering parameters.

To ascertain whether the findings of our simulations are consistent in ‘real world’ scenarios, we re-analyzed two empirical ddRAD datasets using the same combinations of filtering choices. The “empirical low” dataset has low mean read depth and differentiation levels in the range at which our simulations indicate that stringency of call rate threshold is essential. The “empirical high” dataset has high mean read depth and differentiation levels in the range at which our simulations are mostly accurate, regardless of filtering choices. As predicted by our simulations, we see that the two populations in the low differentiation dataset are only detectable when using the more stringent call rate threshold choice, whereas the two populations in the high differentiation dataset are detectable across all combinations of call rate threshold and MAF. Most datasets in our literature review contained poorly differentiated populations; few demonstrated genetic differentiation comparable to that of our most highly differentiated simulations. This may be because high differentiation between conspecifics is biologically rare, or because populations with high levels of differentiation continue to be assessed with less expensive approaches such as microsatellites.

Filtering choices also affected calculations of genetic diversity (He) regardless of inherent levels of differentiation or read depth. At a mean depth of 6, the more stringent call rate threshold and particularly MAF choices produced artificially high He values, while the less stringent filtering produced artificially low He values. This is likely because choosing a stringent MAF value removes loci with very uneven allele frequencies, thereby raising the mean He across loci. However, when mean depth is low and filtering is loose, the retention of obscure, low-quality minor allele calls may undermine evenness and lower the He value. Thus, it is logical that more stringent filtering generally provided results more similar to those of the true genotypes (which were not filtered for MAF). At a mean depth of 25, all filtering choices produced higher values than those calculated from the true genotypes; this is likely because there were much fewer low-quality minor allele calls to be filtered. In this case, looser filtering was likely more consistent with the true genotypes due to the retention of a greater number of high-quality SNP calls. These results demonstrate that estimates of diversity and inferences of population structure depend on filtering and analytical choices, as well as on choice of sequencing depth. Thus, when assessing the validity of choices in filtering thresholds, performing these calculations across combinations of choice parameters would provide useful information and is recommended.

Based on our literature review and analyses of simulated and empirical data, we have framed a recommended set of best practices when analyzing RRS data on wild populations (Box 1). We incorporate and extend previous best practice recommendations for ‘omics data management (Griffin et al., 2017) and adhering to the FAIR principles (Wilkinson et al., 2016). We have shown here that RSS studies often under-report parameter choices, and require improved descriptions of methods (including scripts and metadata) (O’Leary et al., 2018). Raw sequence data should be made accessible to readers in addition to filtered data (O’Leary et al., 2018). This ensures that results are easily reproducible (Gilbert et al., 2012), replicable, and extendible (Griffin et al., 2017).

In this study, we have clarified another aspect of an issue that is increasingly gaining attention: not only do choices of filtering thresholds affect inferences of population structure, but their impact is dependent on the inherent levels of differentiation in the dataset and the mean read depth selected during sequencing. As this body of literature grows, we are optimistic that even more data-sensitive recommendations will emerge, allowing us to continue studying wild populations in efficient and robust ways.

#### Box 1: Recommendations for best practices

- **Use exploratory analyses of varying sequence depths to determine whether actual sequencing depth is sufficient.** It may be useful to first sequence a subset of samples from several of the most geographically separated populations and use these to assess genetic differentiation (F_ST_ and analogues) and actual mean read depth. To avoid over-sequencing, these data could be used in a rarefaction analysis where read depth is reduced to determine the point at which population inferences change.
- **Test a range of filtering parameter choices, present all results, and provide justification for ultimate filtering decisions used in downstream analyses.** For datasets with low differentiation, choice of call rate threshold is critical for accurate detection of population structure. Because global F_ST_ is not strongly affected by filtering parameter choices, it can serve as an indicator of how differentiated the samples in a dataset are, which may inform stringency levels during filtering.
- **Report common measures of differentiation such as global and pairwise F_ST_.** Although 85% of studies in our literature review reported pairwise F_ST_ values, only 28% reported global F_ST_. In addition to providing guidance as to what filtering parameter ranges are appropriate, these measures help researchers gauge how differentiated the populations in a dataset are relative to those in other studies.
- **Report the sample sizes and measures of differentiation and diversity of genetic groups determined by STRUCTURE/PCA, not just those of the groups pre-determined by sampling locations.** If the geographic patterns or number of genetic clusters does not match the geographic patterns or number of populations from which collections were conducted, providing statistics on the actual genetic clusters may be more biologically meaningful and useful to readers.
- In accordance with the **FAIR principles**(Wilkinson et al., 2016):

- **Upload raw reads, not just downstream files in the analysis.** From the 73% of studies that made their data publicly available, only 63% of those uploaded raw reads (the rest uploaded downstream files only, such as VCF files or structure inputs). Making these data available assists other researchers who may want to replicate analyses or to re-analyse data for other purposes; raw reads are likely more useful than are downstream files.
- **Provide accession numbers in manuscripts, and ensure that the accession numbers are correct.** In our literature review, we found that 18% of studies did not provide an accession number, and another 5% provided an accession number, but no files were available at that accession.
- **Provide clear and accurate metadata.** Accurate metadata are critical to re-analyses. Be sure that every sample is accounted for (even samples/files that are removed from the analysis, particularly if they are still uploaded). Provide coordinates to collection sites, as these are key to some population genetic analyses.

## Data Accessibility

Data is available at: https://github.com/m-richardson/Selechnik_et_al_2020_importance_of_filtering_params_for_RRS_studies

## Acknowledgements

This work was supported by the Australian Research Council (FL120100074 to RS and DE150101393 to LAR). The authors would like to thank Rachael Ashby for her constructive feedback during the preparation of this manuscript.

