## Supplementary material for "Inherent population structure determines the importance of filtering parameters for reduced representation sequencing analyses": Table S1

Table S1. Details of population history simulated in each of seven GBS datasets to create varying levels of population structure (as illustrated in main text, Figure 1).

| Dataset | Simulated population history |
| --- | --- |
| Dataset #1  (Global F_ST_ ~ 0.0063) | - 500 founders - Population size increase to 1000 over 100 generations - Constant population size for 900 generations before two-way split between population #1 and populations #2-4 (500 individuals each) - Population #1 undergoes constant population size for 2 generations - Constant population size for 5 generations before two-way split between population #2 and populations #3-4 (250 individuals each) - Population #2 undergoes constant population size for 2 generations - Constant population size for 5 generations before two-way split between population #3 and population #4 (125 individuals each) - Population #3 undergoes constant population size for 2 generations - Population #4 undergoes constant population size for 2 generations |
| Dataset #2  (Global F_ST_ ~ 0.0386) | - 500 founders - Population size increase to 1000 over 100 generations - Constant population size for 100 generations before two-way split between population #1 and populations #2-4 (500 individuals each) - Population #1 undergoes constant population size for 20 generations - Constant population size for 15 generations before two-way split between population #2 and populations #3-4 (250 individuals each) - Population #2 undergoes constant population size for 20 generations - Constant population size for 10 generations before two-way split between population #3 and population #4 (125 individuals each) - Population #3 undergoes constant population size for 20 generations - Population #4 undergoes constant population size for 20 generations |
| Dataset #3  (Global F_ST_ ~ 0.0856) | - 500 founders - Population size increase to 1000 over 100 generations - Two-way split between population #1 and populations #2-4 (500 individuals each) - Population #1 undergoes constant population size for 50 generations - Constant population size for 15 generations before two-way split between population #2 and populations #3-4 (250 individuals each) - Population #2 undergoes constant population size for 50 generations - Constant population size for 10 generations before two-way split between population #3 and population #4 (125 individuals each) - Population #3 undergoes constant population size for 50 generations - Population #4 undergoes constant population size for 50 generations |
| Dataset #4  (Global F_ST_ ~ 0.1576) | - 500 founders - Population size increase to 1000 over 100 generations - Population size decrease to 800 over 20 generations - Population size increase to 1000 over 20 generations - Two-way split between population #1 and populations #2-4 (500 individuals each) - Population #1 undergoes constant population size for 100 generations - Constant population size for 15 generations before two-way split between population #2 and populations #3-4 (250 individuals each) - Population #2 undergoes constant population size for 100 generations - Constant population size for 10 generations before two-way split between population #3 and population #4 (125 individuals each) - Population #3 undergoes constant population size for 100 generations - Population #4 undergoes constant population size for 100 generations |
| Dataset #5  (Global F_ST_ ~ 0.3352) | - 500 founders - Population size increase to 1000 over 100 generations - Population size decrease to 400 over 30 generations - Population size increase to 1000 over 80 generations - Two-way split between population #1 and populations #2-4 (500 individuals each) - Population #1 undergoes constant population size for 200 generations - Constant population size for 15 generations before two-way split between population #2 and populations #3-4 (250 individuals each) - Population #2 undergoes constant population size for 200 generations - Constant population size for 10 generations before two-way split between population #3 and population #4 (125 individuals each) - Population #3 undergoes constant population size for 200 generations - Population #4 undergoes constant population size for 200 generations |
| Dataset #6  (Global F_ST_ ~ 0.6637) | - 500 founders - Population size increase to 1000 over 100 generations - Population size decrease to 200 over 40 generations - Population size increase to 1000 over 120 generations - Two-way split between population #1 and populations #2-4 (500 individuals each) - Population #1 undergoes constant population size for 500 generations - Constant population size for 20 generations before two-way split between population #2 and populations #3-4 (250 individuals each) - Population #2 undergoes constant population size for 500 generations - Constant population size for 15 generations before two-way split between population #3 and population #4 (125 individuals each) - Population #3 undergoes constant population size for 500 generations - Population #4 undergoes constant population size for 500 generations |
| Dataset #7  (Global F_ST_ ~ 0.8103) | - 500 founders - Population size increase to 1000 over 100 generations - Population size decrease to 100 over 50 generations - Population size increase to 1000 over 150 generations - Two-way split between population #1 and populations #2-4 (500 individuals each) - Population #1 undergoes constant population size for 900 generations - Constant population size for 20 generations before two-way split between population #2 and populations #3-4 (250 individuals each) - Population #2 undergoes constant population size for 900 generations - Constant population size for 15 generations before two-way split between population #3 and population #4 (125 individuals each) - Population #3 undergoes constant population size for 900 generations - Population #4 undergoes constant population size for 900 generations |

Table S2. Global and pairwise F_ST_ values with confidence intervals (CI) between two populations, obtained using four different combinations of parameter choices (minimum minor allele frequency=MAF and call rate threshold), on two empirical ddRAD datasets with varying levels of population structure.

| Trumbo et al. (2016) Global and Pairwise F_ST_ (lower limit, upper limit) | | |
| --- | --- | --- |
| Parameter choices | Global | QLD vs NT |
| MAF=0.05, call rate=0.5 | 0.042 | 0.093 (0.091, 0.095) |
| MAF=0.05, call rate=0.2 | 0.036 | 0.097 (0.093, 0.10) |
| MAF=0.01, call rate=0.5 | 0.041 | 0.091 (0.089, 0.094) |
| MAF=0.01, call rate=0.2 | 0.035 | 0.095 (0.092, 0.099) |
| Bell et al. (2015) Global and Pairwise F_ST_ (lower limit, upper limit) | | |
| Parameter choices | Global | Príncipe vs São Tomé |
| MAF=0.05, call rate=0.5 | 0.24 | 0.48 (0.47, 0.49) |
| MAF=0.05, call rate=0.2 | 0.22 | 0.50 (0.49, 0.51) |
| MAF=0.01, call rate=0.5 | 0.21 | 0.43 (0.42, 0.44) |
| MAF=0.01, call rate=0.2 | 0.18 | 0.44 (0.43, 0.45) |

Table S3. Expected heterozygosity (He) of each of two populations, obtained using four different combinations of parameter choices (minimum minor allele frequency=MAF and call rate threshold), on two empirical ddRAD datasets with varying levels of population structure.

| Trumbo et al. (2016) | | |
| --- | --- | --- |
| Parameter choice combination | He QLD | He NT |
| MAF=0.05, call rate=0.5 | 0.50 | 0.20 |
| MAF=0.05, call rate=0.2 | 0.50 | 0.33 |
| MAF=0.01, call rate=0.5 | 0.35 | 0.27 |
| MAF=0.01, call rate=0.2 | 0.47 | 0.28 |
| Bell et al. (2015) | | |
| Parameter choice combination | He Príncipe | He São Tomé |
| MAF=0.05, call rate=0.5 | 0.24 | 0.37 |
| MAF=0.05, call rate=0.2 | 0.21 | 0.50 |
| MAF=0.01, call rate=0.5 | 0.22 | 0.25 |
| MAF=0.01, call rate=0.2 | 0.19 | 0.40 |

(A)

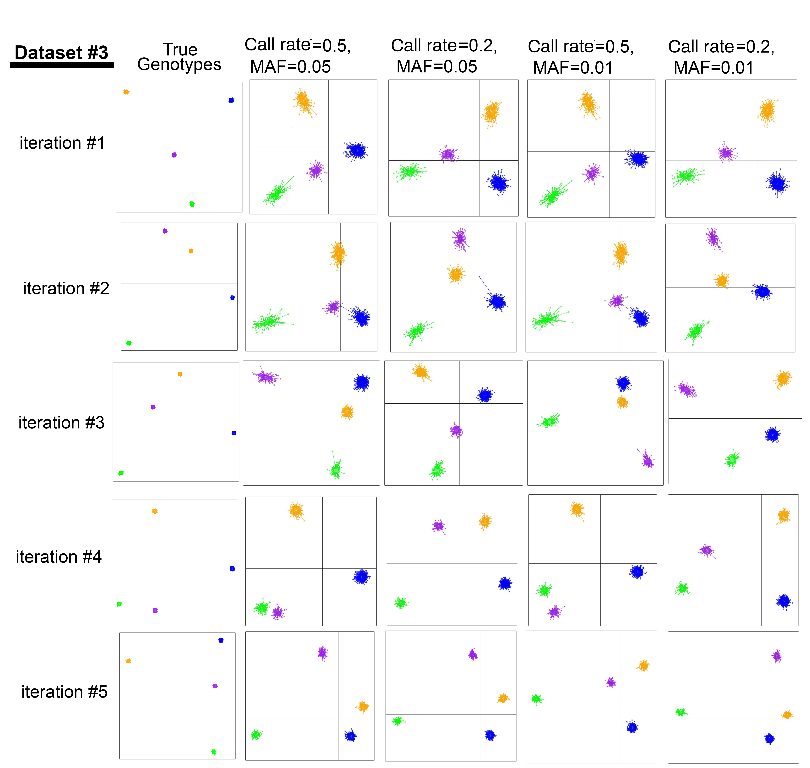

(B)

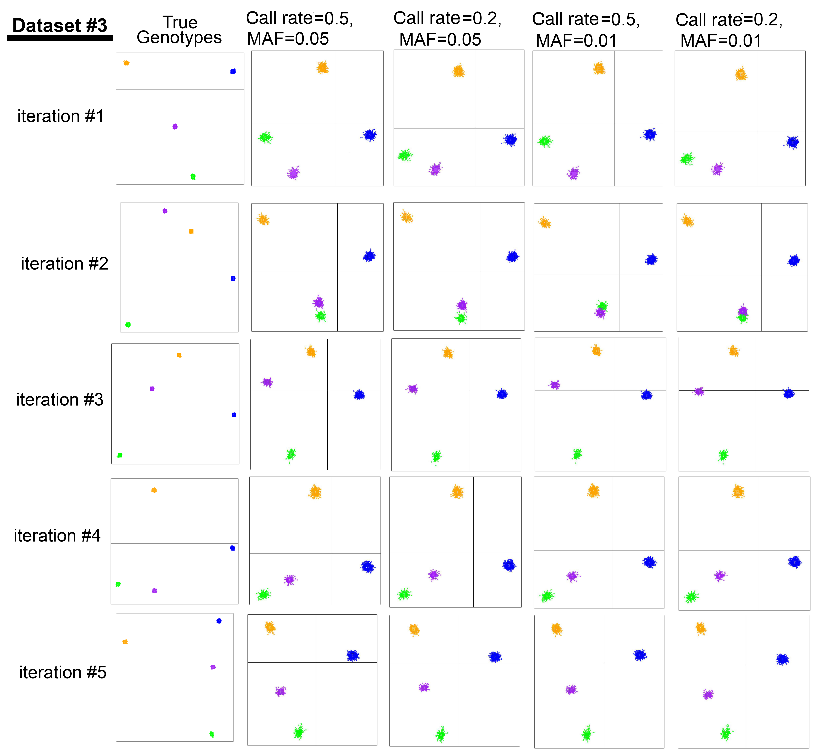

Figure S1. Discriminant analysis of principal components (DAPC) plots of four populations in each of five replicates of a simulated GBS dataset (#3) with relatively low levels of differentiation. These were constructed using observed genotypes for every locus in each dataset. PCAs were run four times with different combinations of filtering parameter choices: minimum minor allele frequency (MAF) and call rate threshold. Read depth was also varied; (A) mean depth = 6 and (B) mean depth = 25.

(A)

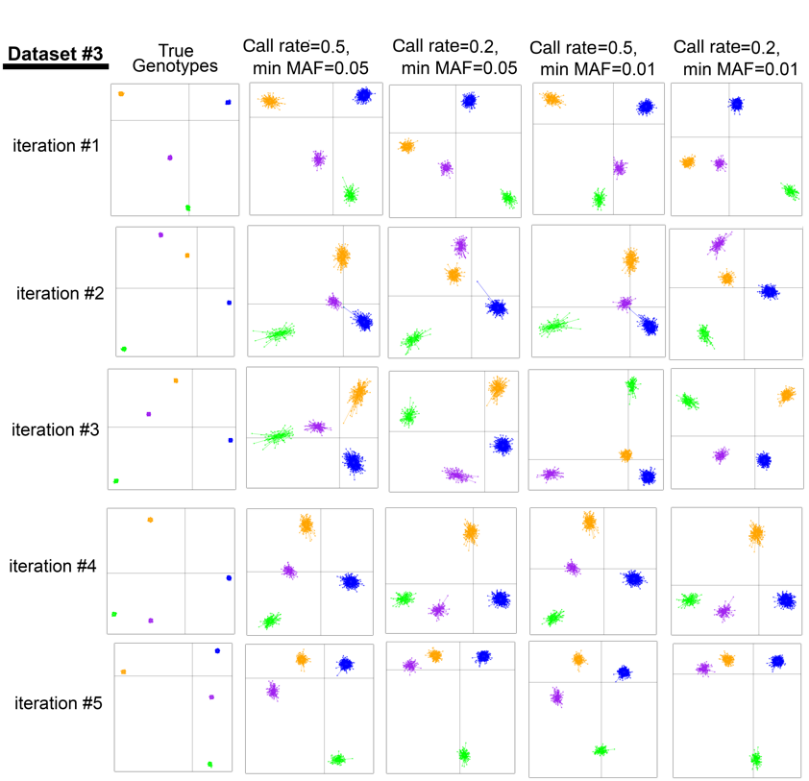

(B)

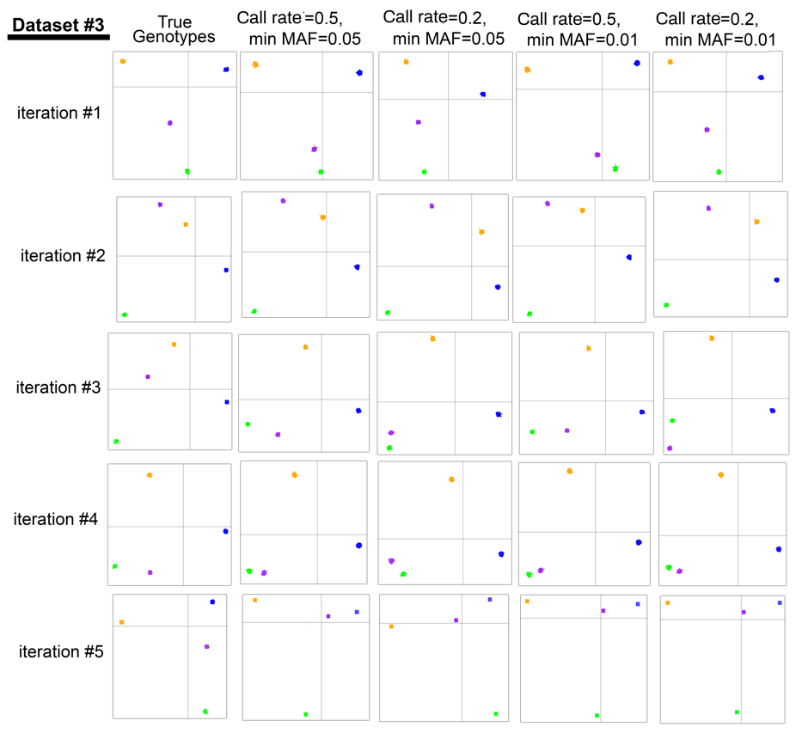

Figure S2. Discriminant analysis of principal components (DAPC) plots of four populations in each of five replicates of a simulated GBS dataset (#3) with relatively low levels of differentiation. These were constructed using allelic dosages (AD) for every locus in each dataset, calculating with genotype probabilities (GP) as follows: AD = 0 × GP homozygous reference + 1 × GP heterozygous + 2 × GP homozygous alternative. PCAs were run four times with different combinations of filtering parameter choices: minimum minor allele frequency (MAF) and call rate threshold. Read depth was also varied; (A) mean depth = 6 and (B) mean depth = 25.

(A)

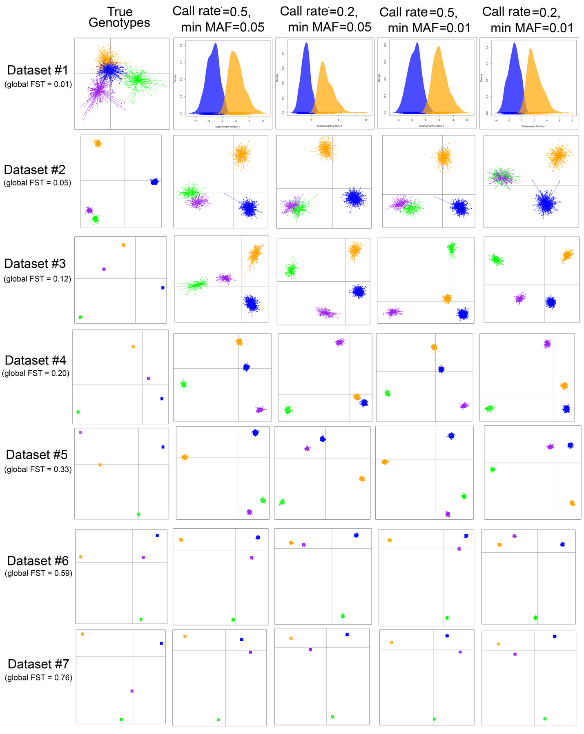

(B)

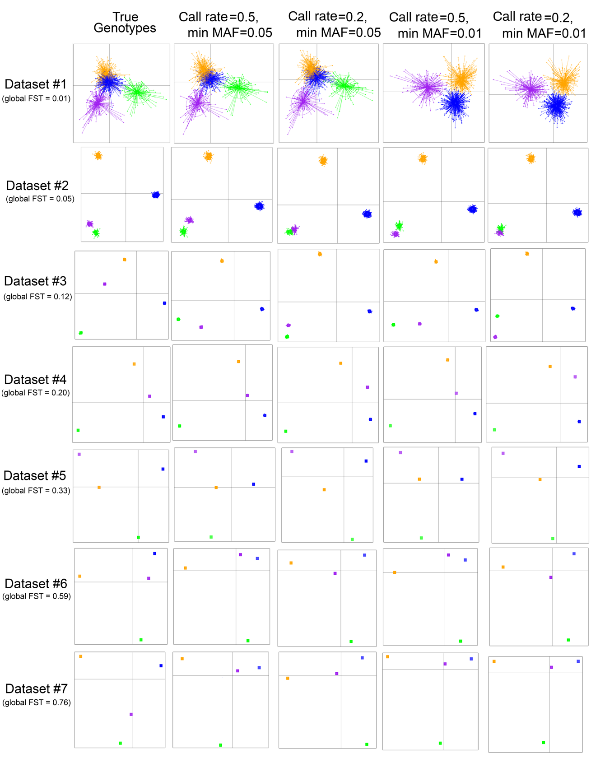

Figure S3. Discriminant analysis of principal components (DAPC) plots of four populations in one replicate of each of seven simulated GBS datasets with varying levels of differentiation. These were constructed using allelic dosages (AD) for every locus in each dataset, calculating with genotype probabilities (GP) as follows: AD = 0 × GP homozygous reference + 1 × GP heterozygous + 2 × GP homozygous alternative. PCAs were run four times with different combinations of filtering parameter choices: minimum minor allele frequency (MAF) and call rate threshold. Read depth was also varied; (A) mean depth = 6 and (B) mean depth = 25. When DAPC detects only two populations (K=2), only a single discriminant function is retained, and densities of individuals on that function are plotted.

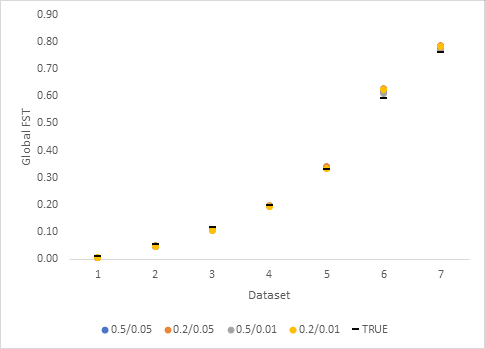

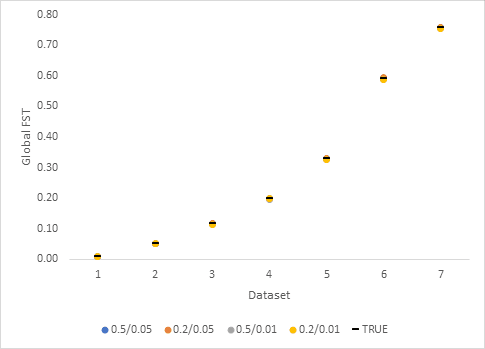

Fig S4A. Global F_ST_ calculations using each parameter combination of call rate and minimum minor allele frequency in relation to Global F_ST_ calculations using the true genotypes (unfiltered; black lines) at a read depth of 6 (top) and a read depth of 25 (bottom).

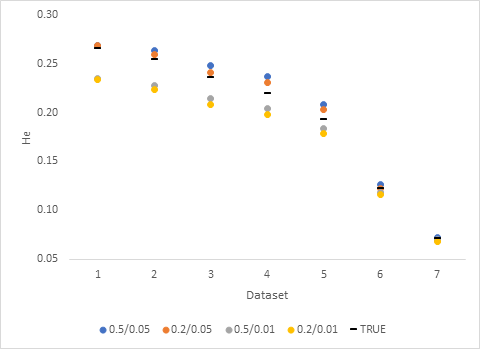

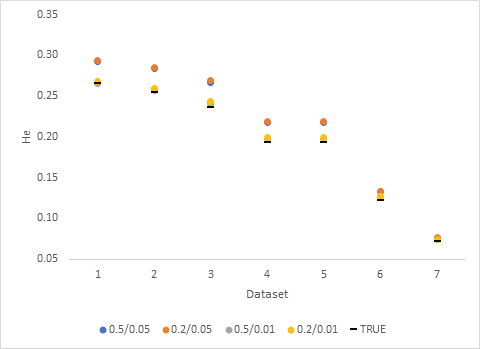

Fig S4B. Expected heterozygosity (He) calculations using each parameter combination of call rate and minimum minor allele frequency in relation to He calculations using the true genotypes (unfiltered; black lines) at a read depth of 6 (top) and a read depth of 25 (bottom).
